# High-density, Identified Cell Recordings from Motor Cortex of Awake Behaving Macaques using 1024-channel SiNAPS-NHP Probes

**DOI:** 10.1101/2025.07.22.665434

**Authors:** G.N. Angotzi, A.M.E. Baker, M. Vincenzi, G. Orban, J.F. Ribeiro, V. Tenorio, L. Berdondini, S. N. Baker

**Author notes:** (co-correspondence and).

## Abstract

**Objective/Background:** Acquiring bioelectric signals from many single neurons in primate brain remains challenging. Chronic implants offer a reasonable channel count (∼100) but sample only a small, fixed region of the cortex. Acutely inserted electrodes can sample from a wider region by making new penetrations each day. The aim of this study was to develop an active dense CMOS probe and experimental procedures to demonstrate acute large-scale single unit recordings from behaving monkeys.

**Methods:** A single-shank CMOS probe was specifically designed for intracortical macaque recordings. The device is based on SiNAPS technology, with additional multiplexing circuits to minimize output lines. Synchronous sampling at 20 kHz/channel from >2k electrode-pixels is achieved with multiple probe systems. Experiments were performed in behaving macaques, achieving multiple day insertions in the motor cortex. Methods were developed to extract spontaneous spiking times of antidromically-identified neurons.

**Results:** The probe (10.7 mm in length, 158 μm in width, 50 μm in thickness) provides a regular array of 1024 electrodes (14 × 14 μm^2^) arranged in 4 columns with an interelectrode pitch of 30 μm. Dural penetration is eased by a small pilot hole; the optimized insertion procedure allows recordings from many different sites. Some cells were identified by antidromic activation as pyramidal tract neurons, which project to the spinal cord.

**Conclusion:** This probe configuration can reach the anterior bank of the central sulcus, which contains many corticospinal cells that connect directly to motoneurons.

**Significance:** This study is an important advance in the toolkit of primate neurophysiology.

## I. Introduction

**N**eural recording technologies for large-scale, single-cell resolution in non-human primates (NHPs), particularly macaques, have seen significant advancements in recent years. While traditional microwire arrays have enabled pioneering long-term studies with recordings from >1,000 isolated units[1], achieving such yields using low-channel-count implants distributed across different brain areas typically requires many recording sessions. In contrast, recent CMOS-based high-density neural probes can achieve comparable or even higher yields per recording session. Moreover, using multiprobe systems enables simultaneous monitoring across multiple brain regions, and can further enhance overall single-unit yield. This supports fine-grained electrophysiological and connectivity studies within localized and distributed brain circuits.

Originally developed for rodent studies, Neuropixels probes have recently been adapted for use in NHPs, enabling high-density, large-scale recordings with single-neuron resolution. The Neuropixels 1.0 [2] and Neuropixels 1.0-NHP [3] allow recordings from 384 channels selected from a larger pool of electrode sites available on the implantable shank, i.e. 960 sites in Neuropixels 1.0, and 2,496 or 4,416 sites in the NHP version, depending on shank length (25 mm or 45 mm). These probes have permitted recordings of over 1,000 neurons using multiple shanks and are now increasingly used in cognitive and sensory neuroscience studies in macaques. A notable application is the Triple-N dataset [4], which used Neuropixels probes to record from 27 subregions of the inferotemporal cortex while monkeys passively viewed 1,000 natural scene images. Such a dataset provides a rich resource for studying visual processing and cross-species comparisons with human data.

Another recent device is the NeuroScroll probe [5], a 1024-channel device with shank lengths ranging from 10 to 90 mm. It has demonstrated reliable chronic recordings in NHPs for up to 105 weeks. However, unlike Neuropixels, NeuroScroll requires external CMOS amplifiers and signal conditioning circuits, necessitating high-density interconnects (up to 179 channels/mm^2^). This increases assembly complexity and introduces experimental handling challenges.

Despite these advances, further progress is needed to enable the reliable acquisition of bioelectrical signals from large, distributed populations of individually identified neurons. This is essential for understanding the implementation and execution of brain functions. For example, motor cortical areas compute the sequence of activation required to produce a given movement. Multiple single unit recordings from these regions have great potential to discover the nature of these computations [6]. Important insights can be obtained by identifying which cells relay the cortical output to subcortical centers. The most prominent motor cortical output neurons are the corticospinal cells. These project long axons through the pyramidal tract at the medulla, and thence on to the spinal cord where they synapse with spinal cord interneurons and, uniquely in primates, to the motoneurons which ultimately relay all motor commands to muscles [7].

Interestingly, pyramidal tract neurons (PTNs) can be straightforwardly identified by antidromic stimulation [8]. This approach involves implanting stimulating electrodes in the pyramidal tract (PT) at the medulla. The placement of the stimulation electrode is performed while monitoring an epidural recording over the motor cortex; it is fixed at the location with the lowest threshold to elicit an antidromic field potential. In subsequent recording sessions, PTNs will respond to PT stimulation with an antidromic spike.

Several criteria can be used to confirm that an evoked spike is antidromic. It should have a low temporal jitter – typically less than 0.1 ms. Larger jitters suggest a synaptic response, produced from collaterals of unrecorded PTNs activating the recorded cell. Secondly, there should be a sharp threshold, with an all-or-none single neuron response appearing over a narrow range of stimulus intensity (typically ∼20 μA). Finally, the response should show collision with spontaneous spikes. Because an action potential leaves in its wake a region of refractory axon, when spontaneous and antidromic spikes meet in the axon between the stimulating and recording electrodes they cannot pass and no antidromic spike is seen in the cortical recording. This occurs when the spontaneous spike occurs over a narrow window around the timing of the PT stimulus.

To advance the current ability to identify a large number of distributed single neurons individually, here we adapted this approach for identifying PTNs and developed a novel high-channel count CMOS probe system based on the SiNAPS technology [9, 10], specifically designed for recordings in the motor cortical areas of awake behaving monkeys (Fig. 1). The realized SiNAPS-NHP probe supports continuous sampling from 1024 electrode channels with a compact I/O interface. It delivers real-time broad-band electrophysiological data at 20 kHz/channel and includes two user-selectable analog channel outputs. The acquisition system supports up to 2048 channels using multiple probe systems. To overcome limitations of chronic implants like the Utah Array (∼100 channels, small, fixed region of the cortex at a fixed depth), we established procedures for repeated acute insertions through the dura. This allows daily targeting of different cortical sites, thus expanding the spatial coverage and flexibility of recordings.

**Fig. 1.**
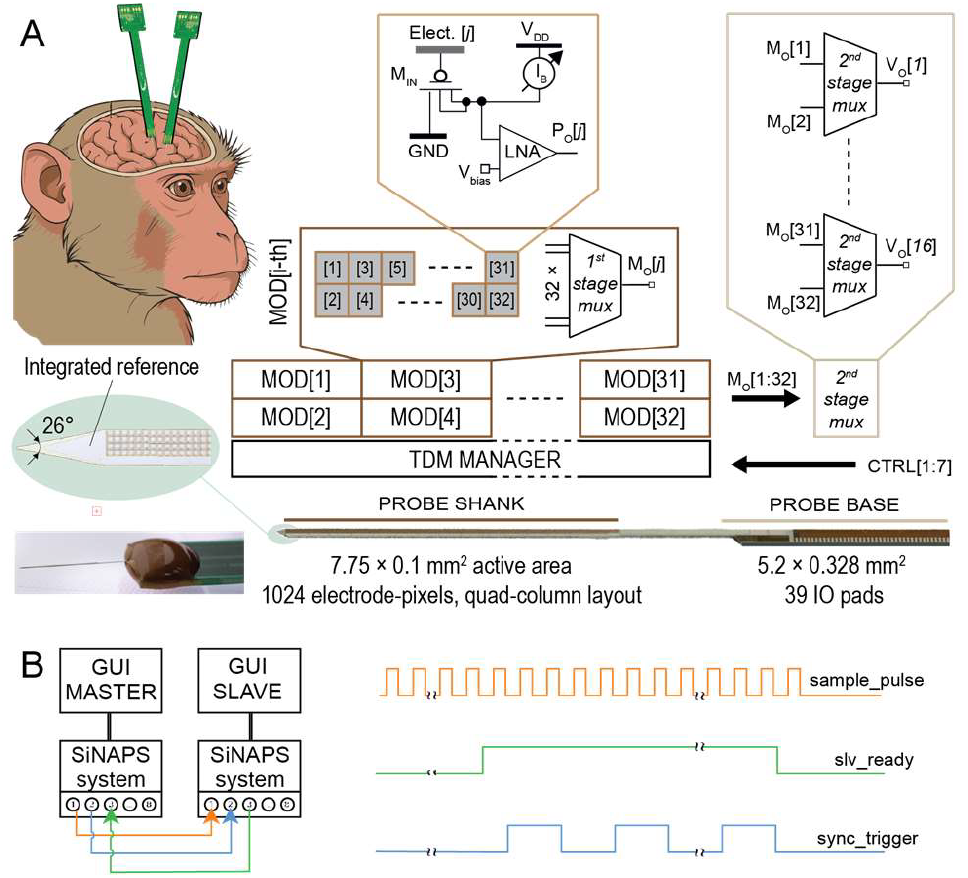
Overview of the 1024 electrode-channels SiNAPS-NHP probe. (A) Illustration of multi-probe recordings in monkeys, with views of the realized probe, and schematic representations of the in-pixel analogue front-end (AFE), on-probe time-division-multiplexing (TDM) and 2^nd^ stage multiplexing (MUX) circuits. (B) Schematics and control signals for synchronous recordings from 2048 electrode-channels using two SiNAPS-NHP probe systems.

## II. Methods

### A. CMOS Probe Design and Realization

A single-shank, high-density CMOS neural probe featuring 1024 electrode channels was developed based on the SiNAPS circuit architecture. This architecture already demonstrated its efficiency for *in vivo* neural recordings [9, 10]. In this design, we modified the electrode-pixel layout, and the architecture was enhanced with additional multiplexing circuits to halve the number of signal outputs, thereby simplifying integration on narrow printed circuit boards (PCBs), see Fig. 1A.

The probe integrates multiple functional modules along a single, elongated shank. Each module contains 32 electrode-pixel sensors, where each pixel comprises a metal electrode pad of 14 × 14 μm^2^ (on the top CMOS metal layer) for neural signal detection, along with underlying signal amplification and conditioning circuits. Consistent with previous SiNAPS designs, each pixel features a DC-coupled analogue front-end (AFE) with a two-stage low-noise amplifier (LNA) and a periodically enabled feedback loop to manage DC offset at the electrode-tissue interface (Fig. 1A). While the amplifier operates primarily in open-loop mode, the autozeroing loop is periodically activated to adjust the bias current of the first stage dynamically, stabilizing the DC operating point.

To streamline data routing, each module employs a 32:1 time-division multiplexing (TDM) scheme, with buffered outputs directed toward the probe base. In this implementation, the main amplifier delivers 40 dB AC gain and incorporates a first-order low-pass filter with a 4 kHz cutoff to minimize aliasing in the TDM stream. Unlike previously reported SiNAPS probes where each module had a dedicated output pad, this design implements a double data rate (DDR) readout scheme at the probe base. The recording area comprises 2 × 16 functional modules arranged along the shank, forming a 4 × 256 electrode-pixel array with a 30 μm pitch, totaling 1024 channels. This layout expands the lateral field of view while maintaining a compact cross-sectional profile suitable for implantation.

Devices were realized in standard 180 nm CMOS technology and post-processed to define the probe shape and electrode material by adapting previously presented processes [11–13]. The microstructuring of the CMOS substrate allowed us to achieve probe shank dimensions of 158 μm x 10.7 mm (active area covered by the electrodes 7.77 × 0.107 mm^2^) and probe base dimensions of 328 μm × 5.2 mm. Additionally, the native 10 × 10 μm^2^ opening on the top CMOS insulator layer – which exposes the top CMOS metal layer (an Al/Cu metal alloy) at each electrode site – was patterned with a 14 × 14 μm^2^ square structured with platinum (Pt) lift-off [11]. By taking advantage of the same lift-off step, an integrated Pt pseudo-reference was patterned next to the electrodes (close-up views in Fig. 2B and 2E). The thickness of the SiNAPS-NHP probes was defined by backside grinding until reaching a 50 μm thick device. Fig. 2 shows SEM pictures of the realized devices. Following experimental validation of the initial prototypes, SiNAPS-NHP probes were realized at wafer-level by Corticale Srl (Genoa, Italy).

**Fig. 2.**
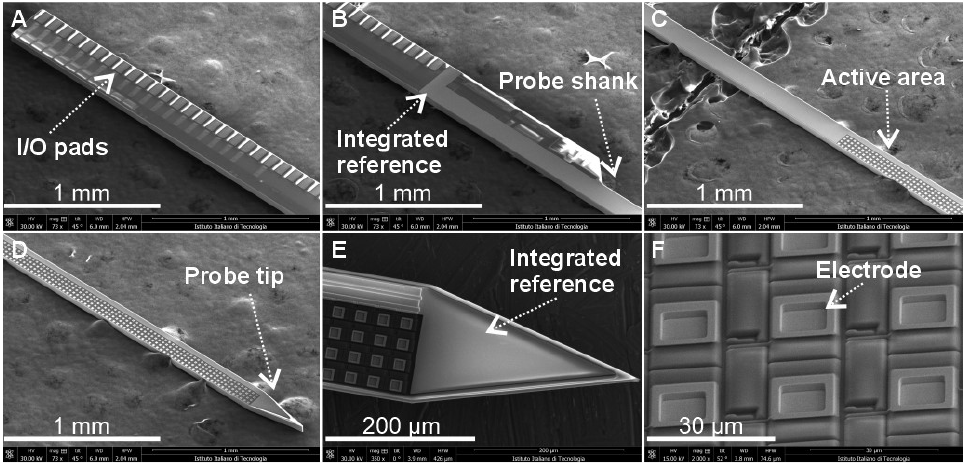
SEM pictures of the realized SiNAPS-NHP active neural probes. The device was realized with a very narrow base to facilitate experimental use. (A) Top of the probe base with the I/O pads. (B) Interface between the probe base and shank with the integrated reference I/O pad. (C) Probe shank with the beginning of the active area. (D) Probe tip. (E) Close-up view of the probe tip with integrated reference. (F) Close-up view of the electrode-pixels.

The probe was then mounted on a narrow PCB (6 mm wide, 1 mm thick, and 5 cm long) and wire bonded. The PCB layout was optimized to allow flexible positioning, enabling access to any point within the exposed craniotomy inside the cranial chamber while also enabling the optional use of the integrated pseudo-reference. Successively, Pt was electrodeposited onto the electrodes to reduce their electrochemical impedance and, consequently, the related noise contribution. Electrodeposition was performed using a potentionstat/galvanostat (PGSTAT204, Autolab Metrohm, Switzerland) with a three-electrode electrochemical cell consisting of an Ag/AgCl reference electrode, a Pt counter electrode and all microelectrodes of the probe connected in parallel as working electrode. Prior to electrodeposition, the Pt microelectrodes were cleaned in 0.5 mol/l (1 N) sulfuric acid (H_2_SO_4_) by cyclic voltammetry (CV) to ensure uniformity. The same cell configuration was then used for Pt electrodeposition by replacing the electrolyte with “Platinum AP + 4G/L Pt” solution (Technic, Italy), and by applying a constant average current of 10 nA per electrode for 1 hour.

### B. Data Acquisition System

The data acquisition (DAQ) platform is built around an FPGA development board based on the Altera Cyclone IV (ZEM4310, OpalKelly Inc., USA), and integrates 12-bit, high-speed, low-power SAR ADCs (MAX11108, Maxim Integrated, USA). It includes power management circuits for stable, low-noise supply from a 5 V input. The FPGA handles time-division multiplexed data readout and digitization (20 kHz, 12-bit per electrode pixel), and runs a customized real-time digital processing unit (DPU) for probe operation. Unlike standard SiNAPS probes, which stream data out on one channel per 32-electrode module, the SiNAPS-NHP probe uses double data rate (DDR) transmission to stream two samples in the same time slot. The DPU also implements a pipelined, real-time second-order IIR filter with a programmable high-pass cutoff (2 Hz or 300 Hz) and a fixed 5 kHz low-pass cutoff (Cortical srl, Italy). For real-time PC interfacing, the FPGA supports high-speed data transfer via CameraLink (raw and processed data) and low-speed command/control via UART.

Two synchronized DAQ systems are used (Fig. 1B) to acquire data from 2048 electrode channels of two SiNAPS-NHP probes. As previously anticipated, for each DAQ, data is acquired frame by frame, with each frame consisting of 1,024 channels read out using a time-division multiplexing (TDM) and double data rate (DDR) scheme. The beginning of each frame is defined by the rising edge of a common *sample_pulse* signal operating at 20 kHz, which is generated by the master system upon being enabled via the master GUI. The master system is enabled first, initiating data acquisition. The same s*ample_pulse* is then sent to the slave system, which responds by asserting the *slv_ready* signal and starting its own acquisition. Since both systems share the same *sample_pulse*, their frame acquisitions are inherently synchronized. Additionally, a *sync_trigger* signal generated by the master and recorded by both systems is used during offline processing to precisely align the two data streams. Finally, each DAQ connects to the corresponding probe via a 1-meter, 50-way cable (P.N. SQCD-025-36.00-TBL-TED-1-S, Samtec, USA). A compact board integrated at the distal side of the cable ensures low-noise delivery of the nominal 1.8 V supply and common reference bias voltages.

### C. Surgical Implant

Experimental recordings have been conducted to date in five macaque monkeys (two male; weight 6.4-12.7 kg). In all cases, animals first underwent an MRI scan under general anesthesia (2-3% sevoflurane inhalation), which allowed the generation of a 3D model of the skull. This was used to produce a digital implant design, based on the annular headpiece first described by [14]. The headpiece was shaped to fit the skull surface, and incorporated a chamber positioned over the primary motor cortex (M1) and dorsal premotor cortex (PMd) (internal dimensions 18×18 mm^2^). The final design was 3D printed in titanium and coated in hydroxyapatite to enhance osteointegration. It was implanted under general anesthesia (2-3% sevoflurane inhalation with 12 μg/kg/hour alfentanil by intravenous infusion), using the system of expanding head bolts to attach the headpiece firmly to the skull also described by [14]. Electromyogram (EMG) electrodes were implanted over 12 muscles in the arm in the same surgery, with wires tunneled subcutaneously to a connector placed on the headpiece. After recovery from the surgery, a craniotomy was opened in the chamber in a further brief surgery under anesthesia to provide daily access for penetrations into the brain. In this surgery, we also implanted two parylene insulated tungsten electrodes (LF501G, Microprobes Inc, Gaithersburg, MD, USA) into the PT at the medulla for stimulation [15, 16].

### D. Pre-Processing and Data Analysis

The recorded data were preprocessed using a custom application developed in MATLAB called SpikeLAB, which provides functionalities such as data visualization in both time and frequency domains, filtering, bad channel characterization, common average referencing (CAR), and single unit sorting. In this work, the recorded data were filtered in the action potential (AP) band and sorting was performed by accessing in SpikeLAB the open-source MATLAB implementation of Kilosort [17]. No manual curation was performed, but a minimum mean firing rate (MFR) threshold was set. Finally, we extracted recording quality indicators, such as i) the signal root-mean-square (RMS) for each channel, ii) the number of masked channels, and iii) the number of units.

### E. Antidromic Identification with SiNAPS-NHP Probes

Once a probe had been inserted into the final location within the cortex, we began by recording responses to PT stimulation, at a fixed intensity chosen to be above threshold for most PTNs (typically 500 μA). PT stimuli were given as biphasic charge-balanced pulses, width 0.1 ms per phase (stimulator model 2100, AM Systems Inc, Sequim, WA, USA). Stimuli were given with a period of 0.5-0.7 s; recordings continued for two minutes. Following this, the PT stimulation was turned off, and recordings made of spontaneous activity during behavioral task performance, typically lasting around one hour. At the end of the session, a further two-minute recording of responses to pyramidal tract (PT) stimulation was made.

To improve the visual distinction between field potentials and antidromic spike responses to PT stimulation, we calculated re-referenced responses *x’* as follows:

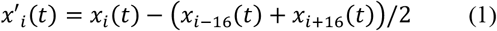

Where *i* is the channel number, *t* is recording time, and *x* is the original recorded data. Because the SiNAPS-NHP probes have four columns of contacts, the offset of 16 channels is equivalent to subtracting the average of the channel four rows above and four rows below from every signal. We expect fields to change slowly with depth, whereas PTN spikes should have a narrowly restricted spatial extent. Spikes should therefore be better visible in *x’* than *x*.

Usually, the problem of spike sorting is carried out in a blind manner – first detecting how many different neurons contribute to the recording and then separating spikes and classifying them. The problem of discriminating identified PTNs is different, since we know *a priori* what the target spike waveform looks like from the antidromic response, in this case across multiple channels. We approached the problem of identifying these spikes by generating a multi-dimensional filter which can be passed over the data to reconstruct the spike train [18, 19]. We model the data by assuming that there is an underlying spike train *S*(*t*), where *S*(*t*)=1 if a spike occurs at time *t* and *S*(*t*)=0 otherwise (*t* is discrete time, indexing the sample number). If the vector *x*_*i*_(*t*) represents the signal from channel *i*=1…*M* of the probe, we extend this by a factor *K* so that the new MK x 1 vector *X*(*t*) contains samples from all *M* channels, over a range of K lags backwards in time from *t*:

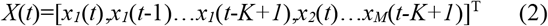

Assuming the occurrence times of a small number of detected spontaneous spikes are *t*_*j*_ (*j*=1..*J*), then the average spike waveform expressed in the extended vector format described above is:

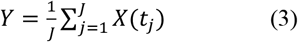

The covariance matrix of the recorded data is defined as:

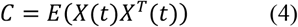

where E(·) is the expectation operator. The spike filter is calculated as:

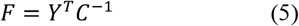

The estimate of the spike train S(t) is provided by:

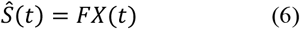

## III. Results

### A. Characterization of SiNAPS-NHP Probes

The impedance of the electrodes (Fig. 3) was measured in NaCl 0.9% using the setup described for Pt electrodeposition. Because individual microelectrodes cannot be individually measured, the measured impedance of a single microelectrode was estimated by multiplying the measured module impedance by the total number of electrodes, thus obtaining an impedance of 858 ± 126 kΩ @ 1 kHz/electrode (n = 5 probes, 1024 electrodes/probe).

**Fig. 3.**
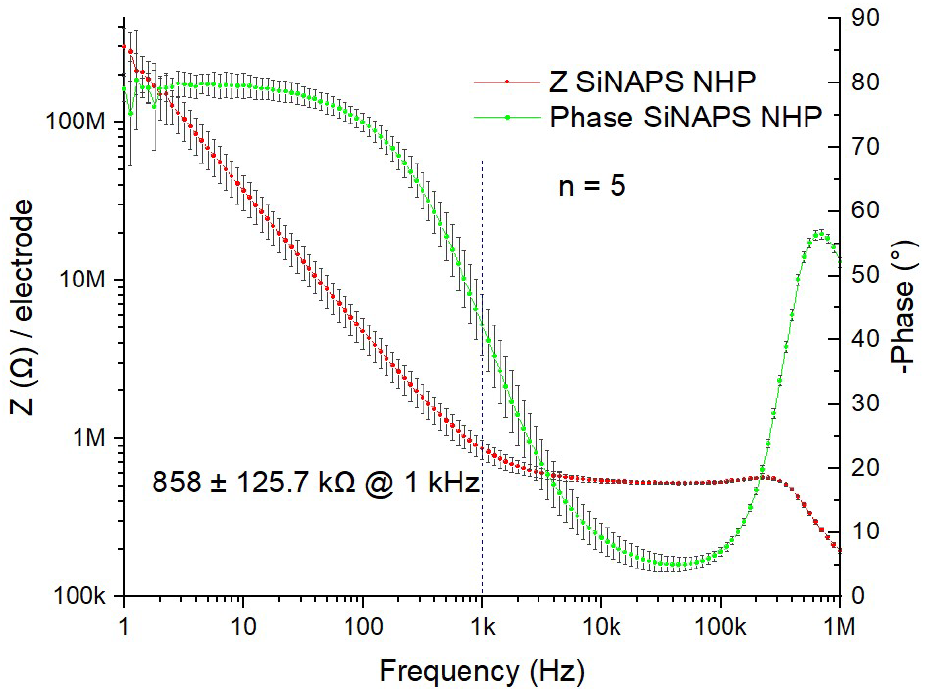
Electrochemical Impedance Spectroscopy (EIS) of the electrodeposited Pt microelectrodes. The mean impedance module at 1 kHz was 858 ± 125.7 kΩ/electrode (n = 5 probes * 1024 electrodes).

The RMS noise was evaluated in saline using a Pt wire to ground the solution (n = 5 probes x 1024 electrodes). The measured noise was: 22.3 ± 6.4 μV_RMS_ for the full-band [0.1 - 5000 Hz], 19.2 ± 7.2 μV_RMS_ for the LFP-band [0.1 – 300 Hz], and 10.5 ± 1.4 μV_RMS_ for the AP-band [300 – 5000 Hz].

### B. Penetration of SiNAPS-NHP Probes into Macaque Cortex

The use of SiNAPS-NHP probes in awake behaving monkeys required us to overcome several technical challenges. The first is the space occupied by the printed circuit board (PCB) which is used to make connections to the silicon probe. When positioned in a recording chamber, the PCB limits how close to the chamber edge the probe can penetrate. For this reason, an important design specification was that the PCB should be as narrow as possible (final width 6 mm). Additionally, we have found that it is essential to visualize the penetration of the probe to avoid breakages. This is aided by a low profile recording chamber design as in [8].

Penetration of a thin probe through the dura mater is difficult. When the dura mater is exposed by a craniotomy inside a recording chamber, there is rapid tissue growth and it becomes thick and tough. This can be prevented using the anti-mitotic agent 5-fluorouracil (5-FU) [8, 20], coupled with meticulous dural care. Tissue growth must be removed daily using fine instruments and visualization under an operating microscope – this is again assisted by the low-profile chamber design. With such care, it is possible to maintain the dura in a thin state for many months, but even with these measures we found that SiNAPS-NHP probes would not penetrate readily into the cortex.

To overcome this issue, the probe was attached to a stereotaxic manipulator mounted on the solid metal plate to which the animal’s head was fixed using the headpiece. This allowed accurate movement of the probe in three dimensions. The probe was angled at 30 degrees to the vertical, so that it penetrated the dura approximately perpendicularly.

In between the probe and the stereotaxic manipulator we used a microdrive (Nan instruments, Nof HaGalil, Israel), which allowed fine control of probe depth. The probe was positioned over a fiducial mark on the side of the chamber, and the manipulator readings noted. We found that placing the probe directly on the metal chamber mark frequently led to breakages. We therefore placed a small piece of hemostatic sponge soaked in saline (Spongostan, Johnson & Johnson) over the mark (Fig. 4B). The soaked sponge was translucent, so that the mark could still be seen. With the sponge in place the moment when the probe contacted the surface could be easily visualized, with less risk of probe breakage.

**Fig. 4.**
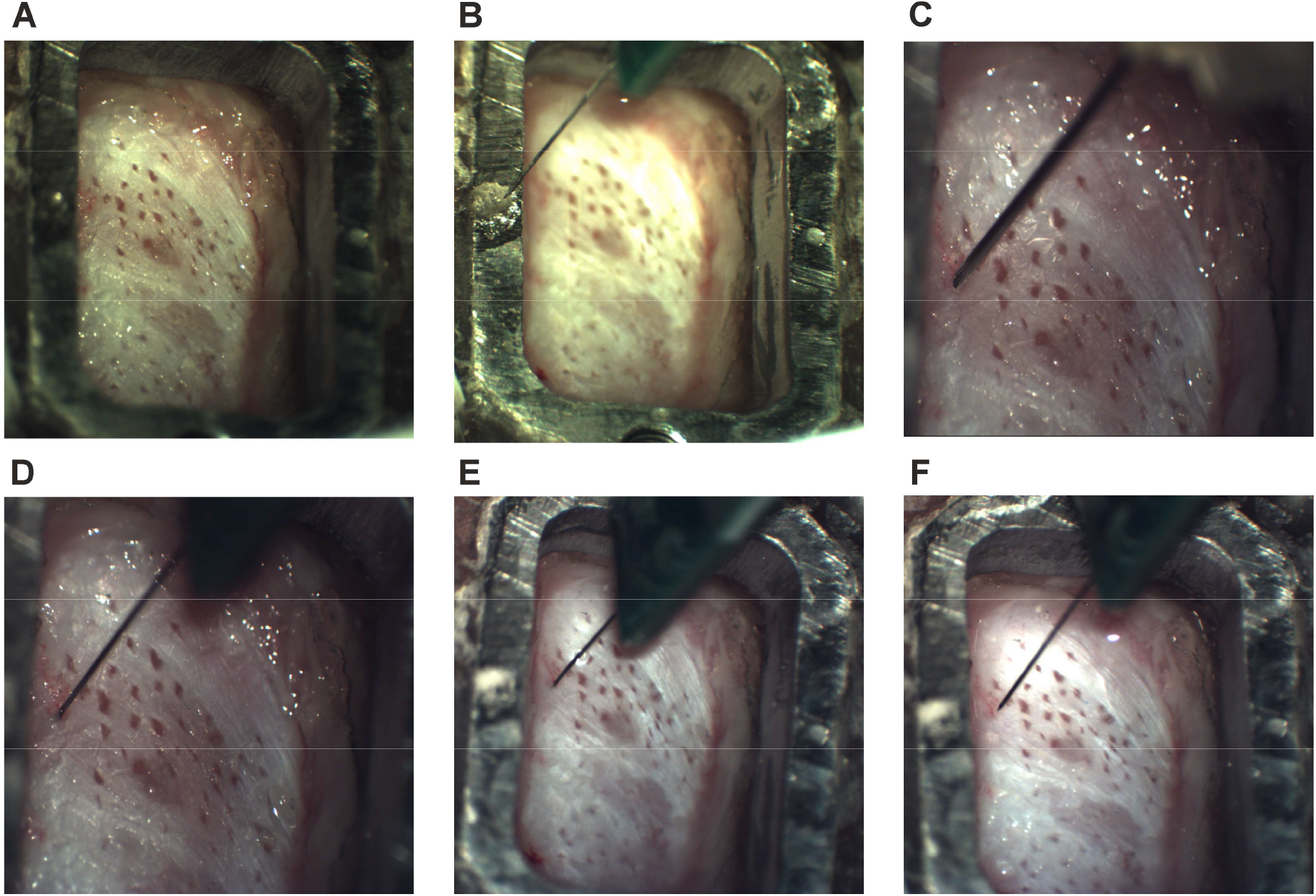
Photographs of recording chamber and sequence of penetration. (A) View of the chamber with exposed dura. (B) SiNAPS-NHP probe located over a piece of Spongostan foam placed over a fiducial mark on the chamber. (C) A 30G needle placed over the intended penetration site over the dura. (D) SiNAPS-NHP probe after penetration through the dura. (E) SiNAPS-NHP probe at final depth for recording. (F) SiNAPS-NHP probe after removal from cortex.

The desired penetration site within the chamber was then calculated relative to the fiducial marker. This used a map of the chamber on which previous penetrations were plotted, and the estimated location of the central sulcus based on the MRI. The manipulator was adjusted to the calculated coordinates, and the probe lowered onto the dura. This location was noted visually, and the probe was raised and moved away. A hand-held 30-gauge needle was then used to make a small puncture at the identified site on the dura (Fig. 4C). Care was taken to angle this needle parallel to the intended probe penetration, and not to pass into the cortex. We found holding the needle by hand more effective than mounting the needle in a manipulator. Using a manipulator, the needle would often dimple the dura, and then suddenly penetrate some distance and damage cells in superficial cortical layers. Such penetrations therefore yielded poor superficial cell recordings. By contrast, the additional visual and proprioceptive feedback available when holding the needle by hand allowed the needle to be rapidly retracted once penetration had been achieved. The photographs in Fig. 4 show the dura covered in small scars caused by previous such penetrations.

The probe was then returned to the intended penetration site, and advanced through the dural hole using the gross height adjustment of the stereotaxic manipulator (Fig. 4D). We found this preferable to using the microdrive, as with manual adjustment of the probe height any failure to penetrate led to clear probe bending; the probe could then be retracted. By contrast, with slow penetration controlled by the microdrive bending was hard to detect and eventually led to breakage. If a probe failed to penetrate, it was removed and a further needle puncture made. We had the impression that this was often due to a failure to pass through not the dura, but the arachnoid underneath. This could also be punctured with a second attempt with the needle and thereby allow penetration. Typically, in successful penetrations the probe was advanced 1-1.5 mm below the surface of the dura.

The probe was then advanced slowly (5-10 μm/s) to the desired recording depth using the microdrive. We found that advancing slowly improved the yield of cells and also reduced drift during recordings. Macaque M1 has a sub-region on the brain surface (‘Old M1’ in the terminology of [21]); when recording here, the probe typically was advanced only 1.5-3 mm (Fig. 4E). When penetrating the part of M1 within the central sulcus (‘New M1’), we could use the full length of the probe and advance 7.7 mm.

Once the probe was at the final depth, recordings were commenced while the monkey performed the behavioral task, which differed depending on the project. Task performance was monitored by single or multi-channel force transducers (3 monkeys), or by high-speed video (2 monkeys). In all cases, a mixture of analog and digital task signals needed to be recorded together with EMG from 12 muscles, as well as the signals from the 1024 contact SiNAPS-NHP probes. This was achieved using an Intan RHD2000 recordings system (Intan Technologies, Los Angeles, CA, USA) running in parallel with the SiNAPS data acquisition device. To synchronize signals for off-line analysis, digital pulses coding task events were sampled by both devices. In some sessions, two SiNAPS-NHP probes were penetrated simultaneously into the cortex. These were connected to two independent SiNAPS data acquisition systems; again, by recording identical task digital events to these as well as the Intan card, the different data streams could be synchronized off-line.

We found that with the workflow described above, probe breakages were rare, and we were able to reuse a single SiNAPS probe for many experimental sessions.

### C. Electrophysiological SiNAPS probe recordings

The data presented in this section were filtered within the 300 Hz to 5 kHz frequency range. Bad channels were masked, and CAR was applied before single units were detected using Kilosort3 with a minimum MFR threshold of 0.1 spikes/s. Out of the 2048 available electrodes, 35 channels (CHs) were masked in Probe1 (P1), and 21 were masked in Probe2 (P2). Figure 5A shows the RMS of the channels computed over a 5 ms time window coinciding with evident neural activity. The higher RMS values correspond to higher spiking activity in the corresponding brain regions. Figure 5B shows the templates of the single units extracted by Kilosort-3 with a firing rate above the threshold, numbering 587 in P1 and 725 in P2. Three single units were selected as a sample for the results presented below and are highlighted in red. To provide a detailed picture of the neuronal activity recorded in regions with higher RMS values, Fig. 5C shows a close-up of the AP signals recorded by the two probes. Each column of electrodes is shown independently, and the signals corresponding to the sample units selected in Fig. 5B are highlighted in black boxes. Finally, Fig. 5D visualizes the templates and autocorrelograms for each sample unit identified in Figs. 5B and 5C.

**Fig. 5.**
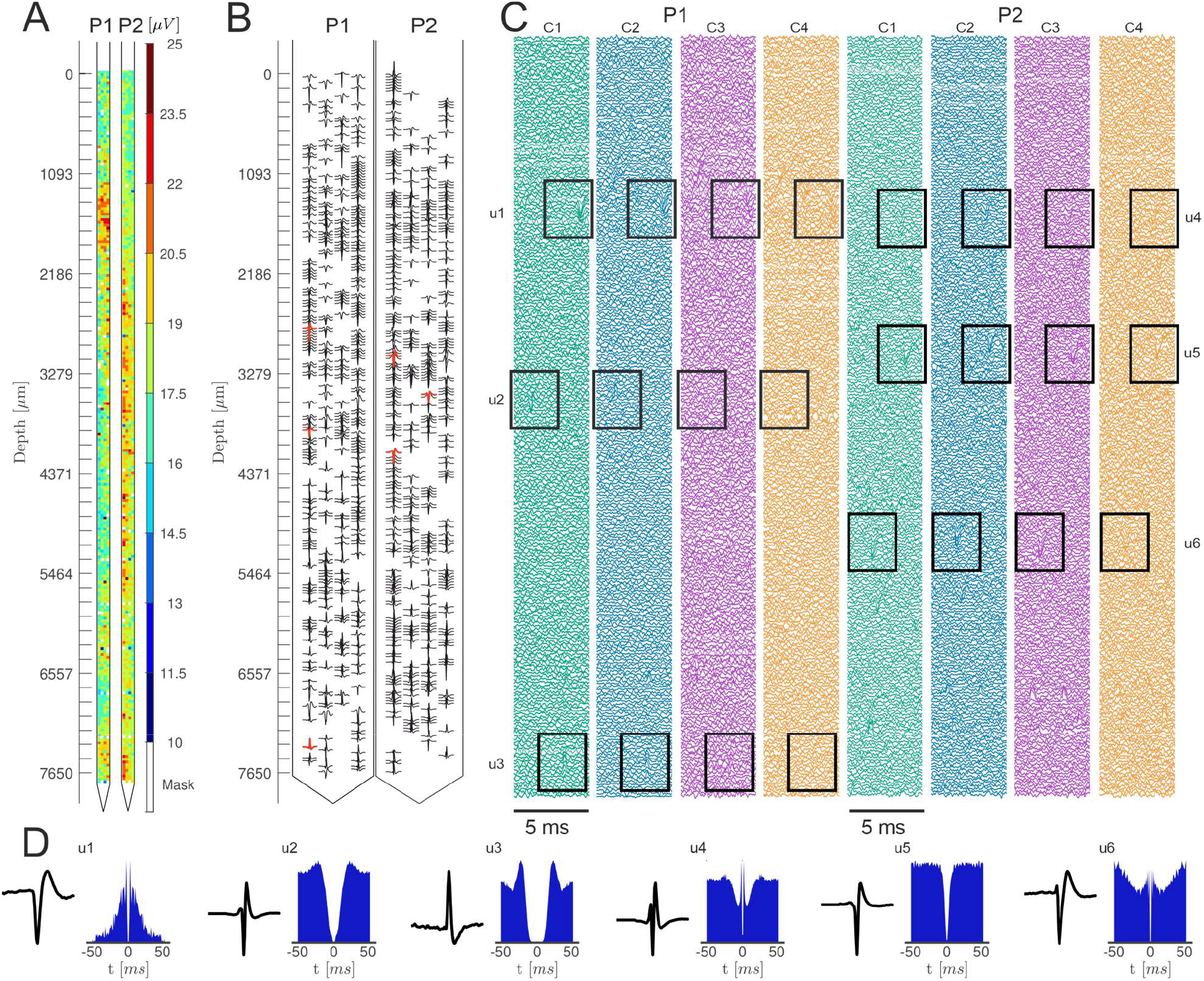
Example of dual SiNAPS NHP probe synchronous recordings for a total of 2048 electrode-channels in the NHP motor cortex. Signals were filtered in the AP band (300 Hz to 5 KHz) and CAR was performed. A total of 35 (over 1024) bad channels (i.e. not at correct working point) were masked in Probe1, and 21 were masked in Probe2. In (A), we represent the channels’ RMS for a time window of 5 ms. In (B), we represent the templates of the single units extracted by Kilosort-3 with an MFR of at least 0.1 spike/s in Probe1 (n=587) and Probe2 (n=725). Three sample units are highlighted in red for each probe.(C) shows a close-up of the signals recorded by the 4 columns (C1-4) of electrodes for each probe. Black boxes highlight the sample units identified in (B). (D) shows the templates and autocorrelograms for each of the sample units displayed in (B) and (C).

### D. Antidromic Identification of Pyramidal Tract Neurons

Offline analysis began by compiling averages of the responses to PT stimulation for each SiNAPS channel and displaying these using a false color map (Fig. 6A) on which the abscissa shows time post-stimulus, and the ordinate channel number. Channel numbers increase with distance from the probe tip, so that the y axis approximately corresponds to the recording location. The large stimulus artifact is immediately apparent (‘art’, Fig. 6A); this is followed by a field potential response (‘field’, Fig. 6A). It is difficult to distinguish any antidromic PTN spikes from this field.

**Fig. 6.**
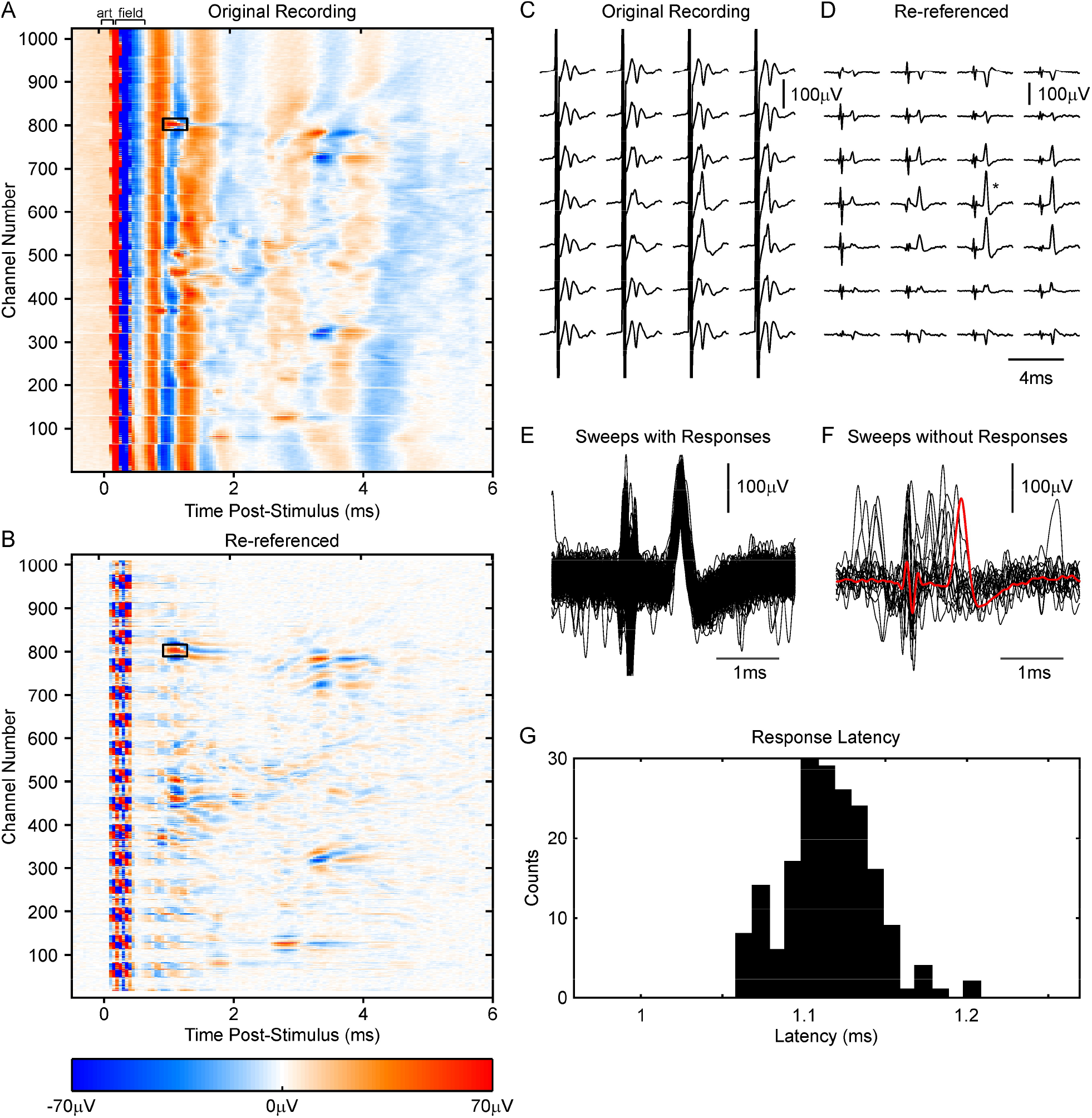
Recording with a SiNAPS-NHP probe of antidromic responses evoked by pyramidal tract (PT) stimulation. A, false color map showing the average response of each channel of a 1024-channel SiNAPS-NHP probe following PT stimulation. An initial stimulus artifact (‘art’) is followed by a field potential (‘field’). Superimposed on this field on some channels are antidromic spike responses. Low channel numbers are deeper within the tissue. B, sweeps as (A), but re-referenced as described in the text. The stimulus artifact was reduced in duration, and the field abolished, allowing antidromic spikes to be more easily visualized. (A) and (B) are shown on the same color scale. C, D, average responses following PT stimulation for the channels indicated by the black box in (A, B). Sweeps are arranged according to the relative geometrical arrangement of the channels. (C) shows original recorded data as in (A), (D) shows re-referenced data as in (B). E, overlain single sweeps for the channel marked by * in (D), for the 187/202 sweeps with a response around 1.1 ms after the stimulus. F, as (E), but for the 15/202 sweeps with no response around 1.1 ms. Overlain red trace shows the average of responses in (E) for comparison. G, histogram of the latency of the spike peak for responses in (E).

Figure 6B shows the average of the re-referenced signal, in which potentials with shallow spatial gradient are removed. Several responses are revealed in this display. One, marked by a black box, is illustrated in more detail in the rest of Fig. 6. While field and spikes are confusingly superimposed in the average of the original data across channels (Fig. 6C), the spike is revealed clearly in the re-referenced data (Fig. 6D).

Fig. 6CD shows average responses; Fig. 6EF shows superimposed single sweeps for the single channel with the largest putative spike response (channel marked by * in Fig. 6D). For this panel, the recording has been reconstructed by convolving the data sampled at 20 kHz with a sinc function [the Nyquist-Shannon theorem, 22]. This provides a high-resolution view of the spike. In most sweeps, there was a clear response (Fig. 6E); however, on 15/202 sweeps, there was no response at the usual latency (compare traces in Fig. 6F with the overlain red trace, which shows the average of the responses from Fig. 6E). In these sweeps, a spontaneous spike occurred before the antidromic latency, leading to collision of the antidromic response. Figure 6G presents a histogram of the spike peak latency for the responses of Fig. 6E. There was extremely low jitter (standard deviation only 27 μs), consistent with an antidromic response.

To detect the spike times of spontaneous spikes produced by the PTN, we first detected the times of a small number of spontaneous spikes at the start of the recording using threshold crossing on the channel with the largest spike in the antidromic response. We selected waveforms similar in shape and size to the antidromic spike by requiring that they pass through voltage windows set manually. Figure 7A presents an average of the antidromic (red) and spontaneous (black) spikes, for the same cell as illustrated in Fig. 6. The close similarity seen in Fig. 7A, coupled with the observed collisions between spontaneous and antidromic spikes (Fig. 6F), provides a high degree of confidence that antidromic and spontaneous spikes come from one and the same neuron. Figure 7B shows a brief section of recording from the probe channel with the largest spontaneous spike for the cell shown in

**Fig. 7.**
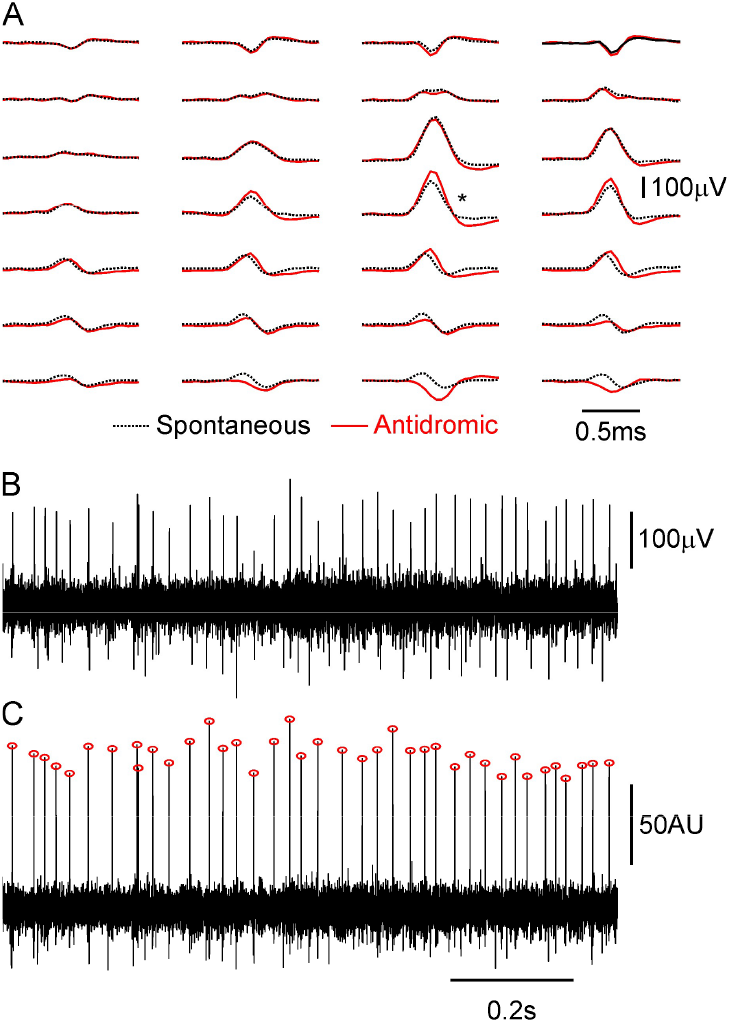
Extracting spontaneous firing of antidromically identified pyramidal tract neuron. (A) Overlain averages of the antidromic spike waveform of the cell shown in Fig. 6 (red), and the spike waveform of a spontaneously active cell (black). The close similarity across channels provides confidence that these are generated by the same neuron. (B) example 1s-long recording from the channel with the largest spike (* in A). (C) example pulse waveform extracted with the multi-channel spike filter. Detected peaks are marked with red circles.

Fig. 7A. Figure 7C shows the reconstructed pulse train *Ŝ*(*t*); peaks in this waveform were detected (MATLAB findpeaks function) and are marked by red circles. The pulse train has a signal to noise ratio (SNR) 2.2x higher than the single channel recording (SNR calculated as spike height divided by noise SD, 13.9 *vs* 6.3). The improvement in SNR obtained by combining multiple channels together optimally has been described previously [23].

### E. Correcting for Electrode Drift

Extracting the occurrence times of spontaneous PTN spikes is straightforward using the approach described above if recordings are stable. However, when making recordings with SiNAPS-NHP probes lasting around one hour in awake behaving monkeys, we frequently observed drift, where the signals moved vertically up the probe. This probably corresponded to the brain tissue adjacent to the probe relaxing gradually after the mechanical stress of driving the probe down. An example of this is shown in Fig. 8. A PTN was clearly visible in the antidromic responses gathered at the start of the experiment (Fig. 8A); one hour later the same cell was also clearly visible (based on its antidromic latency and waveshape across channels), but it had shifted around five rows higher along the probe (Fig. 8B).

**Fig. 8.**
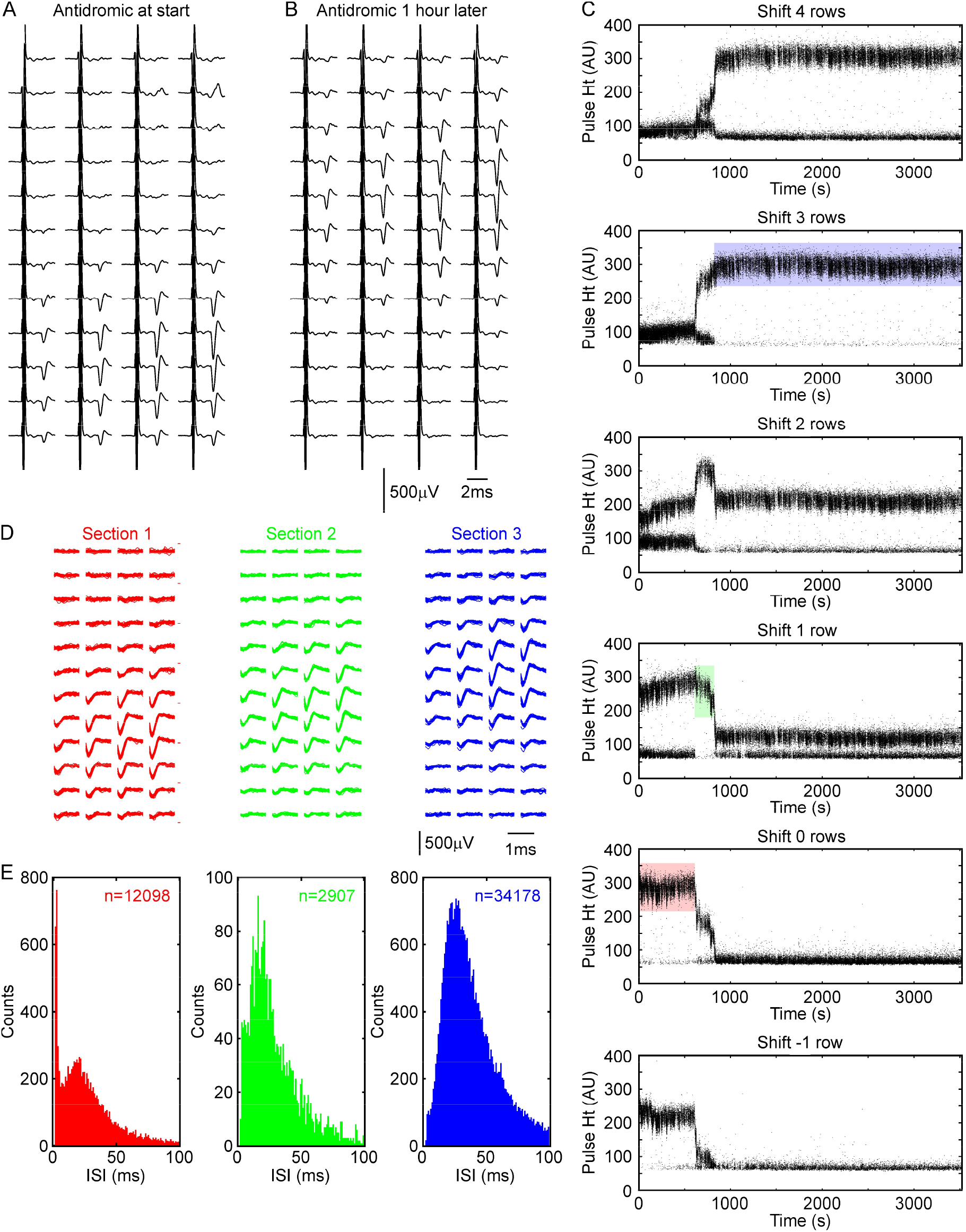
Detecting and tracking drift over a long recording. Average antidromic responses recorded (A) at the start of a session, and (B) at the end one hour later. Note the clear drift in the channel on which the antidromic spike can be seen. (C) Height of peaks in the pulse waveform (corresponding to red circles in Fig. 7) for the entire recorded period. Each plot shows the results when applying the spike filter shifted by several rows up (positive shift) or down (negative shift) relative to the original channels on which the antidromic spike was seen. Sudden shifts in pulse height correspond to movements between probe and tissue. The recording was divided into three sections, and pulses selected from no shift (red, section 1), shift of +1 row (green, section 2) and +3 row (blue, section 3). (D) spike waveforms and (E) interspike interval histograms, for the corresponding color-coded sections. Waveform displays for (A, B, D) show the same channel numbers.

To account for such drift, we assumed that the spike waveshape does not change, but that it was merely translated by an integer number of rows up or down the probe. We recalculated a range of filters for different shifts and extracted the corresponding pulse train estimates *Ŝ*(*t*). Figure 8C plots the heights of the peaks of the pulse train for various shifts. At the start of the recording, the pulse train with no shift had peaks which were clearly separated from the noise (Fig. 8C, red box); selecting these heights extracted a clean single unit, with consistent waveform (Fig. 8D, red). The inter-spike interval histogram (Fig. 8E, red) showed a lack of intervals within the first 1 ms, consistent with the neuronal refractory period. At around 611 s into the recording, a sudden change occurred, and the peak height at zero shift reduced. However, peaks remained clearly separated from the noise in the pulse train extracted by shifting one row up the probe (Fig. 8D, green), allowing continued clean discrimination. At 823 s after the recording start, there was another change, which necessitated shifting a further two rows up the probe (Fig. 8E, blue). The inter-spike interval histograms over these three sections were broadly consistent, although the first section showed a narrow peak at an interval of 3 ms reflecting burst firing which was not seen in the other two. This may be caused by damage to the neuron and its dendritic tree shortly after probe penetration, leading to injury discharges; we assume that after the damaged neuron membrane resealed these injury discharges ceased.

The total shift required to discriminate the pulse train from the beginning to the end of the file was three rows, although equally good separation could be achieved with a shift of four rows (Fig. 8C, top). This compares with an estimated five row shift based on antidromic response in Fig. 8B. It is possible that some additional shift occurred between the end of the recording of task-related activity and the start of the second file measuring antidromic responses. In between these two files, the monkey typically makes multiple postural adjustments after the long period of focused task performance. It is therefore highly plausible that there could be an additional movement of tissue relative to probe at this time.

## IV. Conclusion

In this study, we have described the design and manufacture of a new CMOS-based probe which considerably expands the state of the art in recording from the cerebral cortex in awake behaving monkey. We have shown that the high channel count, all of which are sampled, allows large numbers of neurons to be simultaneously recorded across a wide depth. Furthermore, we describe an approach to antidromic identification of a sub-set of the recorded neurons. Knowing which cells project to an output region (in the case of M1, the spinal cord) is critical information in understanding the network computations performed by a cortical area. We suggest that SiNAPS-NHP probes are an important advance in the toolkit of primate neurophysiology.

## Acknowledgment

We thank Terri Jackson and Andrew Atkinson for animal training, Rocio Palacios-O’Connor for veterinary support, Michelle Waddle for surgical theatre nursing, and Norman Charlton for mechanical engineering support. We would like also to thank Corticale Srl (Italy) for having scaled up the fabrication of the SiNAPS 1024 NHP probes and for their support on the firmware.

